# APOE4 Increases Susceptibility to Amyloid, Accelerating Episodic Memory Decline

**DOI:** 10.1101/2024.12.23.630203

**Authors:** Casey R. Vanderlip, Craig E.L. Stark, the Alzheimer’s Disease Neuroimaging Initiative

## Abstract

Apolipoprotein E4 (APOE4) is the strongest genetic risk factor for sporadic Alzheimer’s disease (AD). Individuals with one copy of APOE4 exhibit greater amyloid-beta (Aβ) deposition compared to noncarriers, an effect that is even more pronounced in APOE4 homozygotes. Interestingly, APOE4 carriers not only show more AD pathology but also experience more rapid cognitive decline, particularly in episodic memory. The underlying mechanisms driving this domain-specific vulnerability, however, remain unclear. In this study, we examined whether the accelerated decline in episodic memory among APOE4 carriers is due to increased Aβ deposition or heightened susceptibility to Aβ-related effects. Using data from the Alzheimer’s Disease Research Initiative, we modeled amyloid duration, the estimated number of years an individual has been amyloid-positive, and its impact on cognitive trajectories. Our findings reveal that APOE4 is associated with more rapid episodic memory decline as a function of amyloid duration. This decline was dose-dependent, with APOE4 homozygotes declining more rapidly than heterozygotes, and it was consistently observed across multiple episodic memory tasks and measures. Importantly, this pattern was not observed in other cognitive domains, such as processing speed, executive function, visuospatial skills, language, or crystallized intelligence. These results suggest that cognitive trajectories in AD differ by APOE genotype, with APOE4 conferring increased vulnerability to hippocampal dysfunction early in the disease course. Future research should investigate whether these cognitive differences stem from distinct pathological cascades in APOE4 carriers.

## 1. Introduction

Alzheimer’s Disease is primarily defined by the accumulation of amyloid-beta (Aβ) and tau in the brain, which are considered the core pathologies of the disease. However, debate exists regarding whether cognitive impairment is essential for a clinical diagnosis. Recent guidelines from the Alzheimer’s Association workgroup state that elevated Aβ and tau biomarkers alone are sufficient for an AD diagnosis, even in the absence of cognitive decline. In contrast, the International Working Group emphasizes that both biomarkers and cognitive impairment are necessary, highlighting the clinical aspect of diagnosis. Importantly, while the Alzheimer’s Association workgroup considers Aβ and tau as definitive indicators of AD, they recommend diagnosing individuals only when cognitive impairment is present. Therefore, regardless of criteria, detection of AD-related cognitive impairment is critical for timely diagnosis and intervention. However, this requires a comprehensive understanding of cognitive impairment and whether subtypes of AD may present with different patterns of impairment.

Apolipoprotein E4 (APOE4) is the most significant genetic risk factor for developing sporadic AD. While most individuals carry two copies of APOE3, approximately a quarter of the population are APOE4 carriers, which more than doubles their risk of AD (Genin et al., 2011; Gharbi-Meliani et al., 2021). Furthermore, around 2% of the population has two copies of APOE4, yet this group accounts for a quarter of AD cases. These individuals not only face a higher risk of AD but also develop pathology, such as Aβ and tau deposition, earlier than non-carriers (Fortea et al., 2024; Jansen et al., 2015). APOE4 carriers are more likely to develop dementia and have an earlier mortality rate (Corder et al., 1993; Reiman et al., 2020). This increased likelihood of AD in APOE4 homozygotes has led to the hypothesis that APOE4 homozygosity may represent a genetic form of AD (Fortea et al., 2024).

Aβ deposition is considered an early, if not the earliest, pathology in the development of AD (Jack et al., 2018; Sperling et al., 2011). Research has shown that APOE4 carrier status is associated with increased Aβ deposition, which begins at a younger age compared to APOE3 homozygotes and even earlier in APOE4 homozygotes (Belloy et al., 2019; Morris et al., 2010). By age 80, nearly all APOE4 homozygotes are Aβ positive, and approximately 80% of APOE4 heterozygotes also show Aβ positivity (Fortea et al., 2024). Interestingly, once individuals are Aβ positive, the rate of Aβ accumulation does not differ significantly between APOE genotypes (Betthauser et al., 2022; Lim et al., 2017). Therefore, it is plausible that the association of APOE4 with increased cognitive decline and dementia is due primarily to the earlier onset of disease pathology in these individuals.

While AD associated dementia is associated with global cognitive impairment, not all cognitive domains are equally affected. The earliest cognitive impairments in AD typically involve tasks that engage the hippocampus, such as episodic memory (Gallagher & Koh, 2011; Grande et al., 2021). Research has shown that deficits in episodic memory can serve as biomarkers for AD pathology and predict future cognitive decline in cognitively normal older adults (Berron et al., 2024; Vanderlip, Lee, et al., 2024; Vanderlip, Stark, et al., 2024). Notably, APOE4 carriers tend to experience greater age-related deficits in episodic memory and show faster decline in this domain over time, particularly in individuals with elevated Aβ deposition (Bondi et al., 1995; Eich et al., 2019; Lim et al., 2016; Mormino et al., 2014) This supports the notion that APOE4 carriers may be further along the AD spectrum. However, evidence suggests that this accelerated decline is not observed in other cognitive domains, such as executive function or language, implying that the faster decline in APOE4 carriers may be specific to episodic memory (Lim et al., 2016).

Based on these findings, we propose two potential explanations. First, APOE4 carriers may be further along the Aβ spectrum, with increased pathology leading to more pronounced memory decline. Alternatively, APOE4 may enhance susceptibility to Aβ, meaning that less pathology is required to trigger memory deficits in these individuals. Given that episodic memory appears to be particularly vulnerable in APOE4 carriers, we hypothesize that the brain regions supporting this cognitive domain, such as the hippocampus, may have a heightened susceptibility to Aβ in these individuals.

Extensive research has shown that once individuals reach a certain threshold of Aβ, its accumulation proceeds at a similar rate across different people (Betthauser et al., 2022; Farrell et al., 2021; Insel et al., 2021; Jagust et al., 2021). Consequently, studies have modeled Aβ duration, the number of years an individual has been Aβ positive and used this to examine the time course of other pathological changes and cognitive decline in AD (Cody et al., 2024; Jia et al., 2024; Li et al., 2024). This critical technique allows researchers to investigate when changes in AD begin and helps address whether APOE4 carriers are more susceptible to Aβ deposition. In this study, we investigated the time course and severity of cognitive changes as a function of Aβ positivity and APOE genotype, focusing on the interaction between these two factors in a large multicenter study.

## 2. Methods

Data used in the preparation of this article were obtained from the Alzheimer’s Disease Neuroimaging Initiative (ADNI) database (adni.loni.usc.edu). The ADNI was launched in 2003 as a public-private partnership, led by Principal Investigator Michael W. Weiner, MD. The primary goal of ADNI has been to test whether serial magnetic resonance imaging (MRI), positron emission tomography (PET), other biological markers, and clinical and neuropsychological assessment can be combined to measure the progression of mild cognitive impairment (MCI) and early Alzheimer’s disease (AD).

### 2.1. Participants

Seventeen hundred and one older adults who underwent neuropsychological test and underwent Aβ imaging were included (Table 1). No participants had a history of major neurological or psychiatric disorders, head trauma, or history of drug abuse or dependency.

**Table 1:**
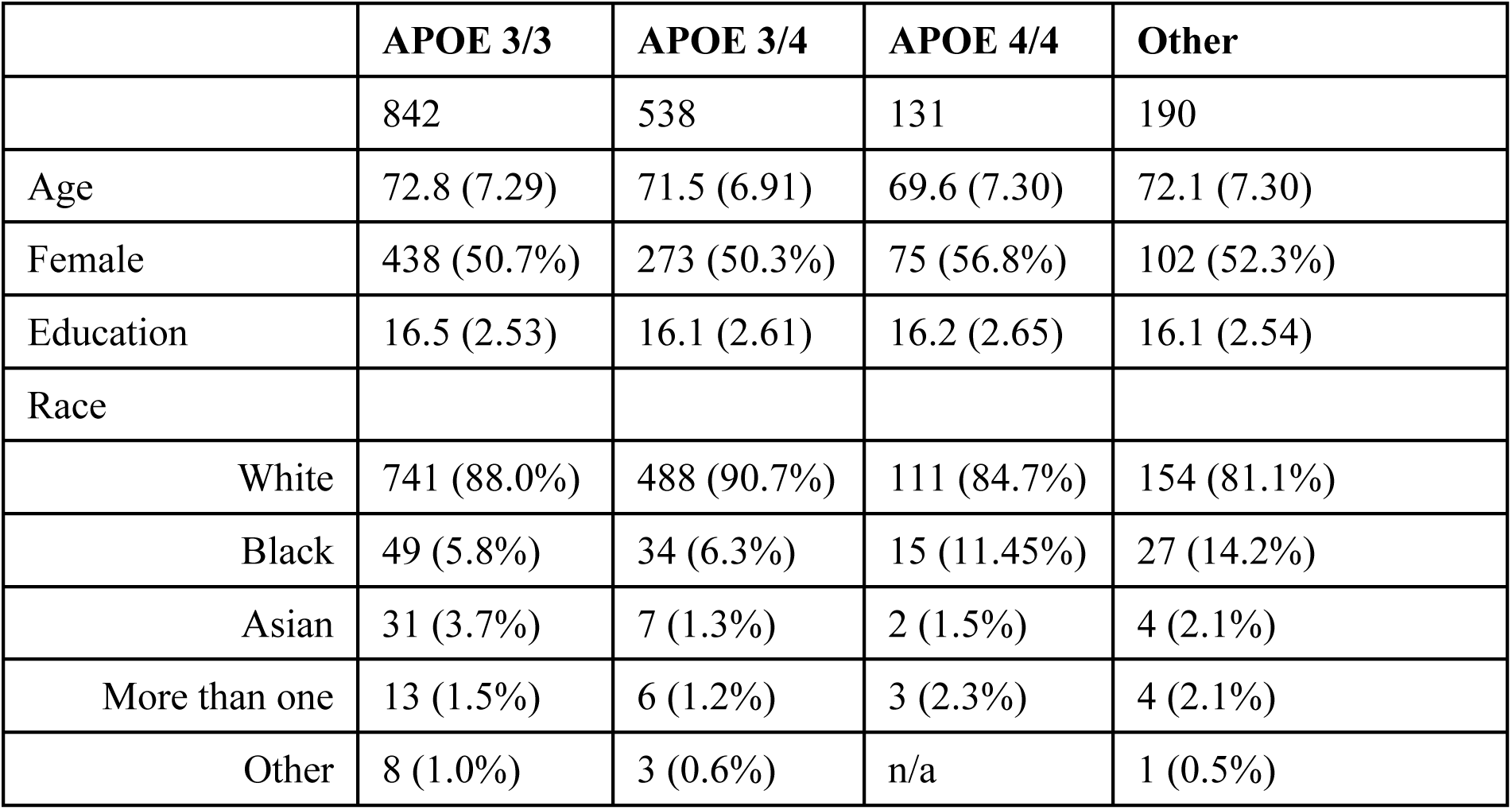
Demographics.

|  | **APOE 3/3** | **APOE 3/4** | **APOE 4/4** | **Other** |
| --- | --- | --- | --- | --- |
|  | 842 | 538 | 131 | 190 |
| Age | 72.8 (7.29) | 71.5 (6.91) | 69.6 (7.30) | 72.1 (7.30) |
| Female | 438 (50.7%) | 273 (50.3%) | 75 (56.8%) | 102 (52.3%) |
| Education | 16.5 (2.53) | 16.1 (2.61) | 16.2 (2.65) | 16.1 (2.54) |
| Race |  |  |  |  |
| White | 741 (88.0%) | 488 (90.7%) | 111 (84.7%) | 154 (81.1%) |
| Black | 49 (5.8%) | 34 (6.3%) | 15 (11.45%) | 27 (14.2%) |
| Asian | 31 (3.7%) | 7 (1.3%) | 2 (1.5%) | 4 (2.1%) |
| More than one | 13 (1.5%) | 6 (1.2%) | 3 (2.3%) | 4 (2.1%) |
| Other | 8 (1.0%) | 3 (0.6%) | n/a | 1 (0.5%) |

### 2.2. Neuropsychological testing

All participants underwent comprehensive neuropsychological testing, which has been described in detail and described briefly, below (Crane et al., 2012). To quantify memory ability, we used performance from the Rey Auditory Verbal Learning Test (RAVLT) and Logical Memory Test. The RAVLT is a list learning paradigm where participants learn a series of 15 words from a list “A” over 5 trials followed by one trial of an interference list “B”. Afterwords, participants recalled words from list “A” immediately and again after a twenty-minute delay (Delayed Recall). Further, after the delayed recall, participants were read a list of 30 words and asked if the word appeared on list “A”. Participants made yes/no judgements and performance on this score was quantified (Recognition). The logical memory test involves an experimenter reading a participant a short story and asking the participant to immediately recall the details of the story and again after a delay of twenty minutes (Delayed Recall). Here we used Delayed Recall from both the RAVLT and Logical Memory test along with Recognition performance on the RAVLT as measures of memory ability.

In addition to the three memory-based measures, we also utilized non-memory tasks including Trail Making Test (TMT), Clock Drawing Task, Boston Naming Test (BNT), and the American National Reading Test (ANART). The TMT was administered to assess processing speed and cognitive flexibility. In TMT-A, participants connected numbered circles (1–25) in sequential order as quickly as possible. In TMT-B, they alternated between numbers (1–13) and letters (A–L) in ascending and alphabetical order. Participants had up to 300 seconds to complete each part and completion times (in seconds) were recorded. Time to complete TMT-B is thought to reflect processing speed, while TMT-B time is used to evaluate cognitive flexibility.

The Clock Drawing task was administered to assess visuospatial and executive function. Participants were instructed to draw a clock face, place the numbers correctly, and set the time to “10 past 11.” Scoring (0–5 scale) was based on the accuracy of the clock’s numbers and hands. The BNT was used to assess language and word retrieval abilities. Participants were shown 60 black-and-white line drawings and asked to name each item. If a participant struggled to name an object, a semantic cue was provided, followed by a phonemic cue if needed. The primary outcome measure was the total number of correct responses, with partial credit given for responses following cues. The ANART was administered to estimate crystalized intelligence. Participants were asked to read aloud a list of 50 irregularly spelled words. The number of pronunciation errors was recorded, with fewer errors indicating higher performance.

### 2.3. PET imaging

All individuals underwent either underwent flobetapir (FBP) (n = 2595) or florbetaben (FBB) (n = 401) imaging to quantify Aβ. Preprocessing and quantification of the data was handled by the ADNI PET core. Aβ centiloid values were provided and used for all analyses. Comprehensive information regarding the PET processing and acquisition techniques is available on the ADNI website at https://adni.loni.usc.edu/wp-content/uploads/2012/10/ADNI3_PET-Tech-Manual_V2.0_20161206.pdf.

### 2.4. Estimating Aβ duration

Previous research has shown that the onset of Aβ deposition varies significantly across individuals. However, after surpassing a specific threshold of Aβ accumulation, individuals tend to follow a consistent progression. These findings have been widely validated and applied to track the progression of other AD-related pathologies (Betthauser et al., 2022; Jia et al., 2024; Li et al., 2024). To estimate the number of years individuals were Aβ positive, we employed Sampled Iterative Local Approximation (SILA) to model Aβ accumulation over time (Betthauser et al., 2022). SILA iteratively sampled centiloid values from the cortical composite ROI to generate a continuous timeline of Aβ burden. We used a positivity threshold of 16.7 centiloids, as this level is considered a critical tipping point Aβ deposition in the ADNI dataset (Farrell et al., 2021; Li et al., 2024; Schindler et al., 2021) and we wanted to identify those at the earliest stages of AD. The estimated years of Aβ positivity for each subject was calculated as the difference between the predicted age of Aβ onset and the age at the time of each neuropsychological assessment.

### 2.5. Statistical Analyses

Data were analyzed using R. Cognitive scores were z-scored to standardize performance, and the direction of TMT and ANART scores was reversed so that lower z-scores indicated poorer performance. We controlled for confounding variables by regressing out age, sex, and education from each cognitive score and analyzing the residuals. Analyses focused on individuals with an APOE genotype of 3/3, 3/4, or 4/4. To assess cognitive changes as a function of years of Aβ positivity, LOESS curves were fitted for each genotype. Bootstrapping (1,000 iterations) was used for confidence intervals and were used to compare genotypes, and AUCs were calculated using the trapezoidal rule. If confidence intervals of the bootstrapped AUCs did not overlap between genotypes, comparisons were considered significant. Two-way ANOVA was conducted to examine interactions between factors, with a significance threshold set at p < 0.05.

## 3. Results

### 3.1. APOE4 carriers exhibit increased, but not faster, Aβ deposition

Extensive research links Aβ deposition with increasing Aβ burden over time. To investigate this, we analyzed the association between age and Aβ centiloid values using LOESS smoothing, stratified by genotype (Figure 1A). We formally compared Aβ deposition across genotypes by calculating the area under the LOESS curves from 1,000 bootstrapped iterations, generating 95% confidence intervals for the AUCs: APOE3/3 [454, 992], APOE3/4 [1408, 2038], and APOE4/4 [2121, 2896]. Significant increases in Aβ AUCs were observed for APOE3/4 and APOE4/4 compared to APOE3/3, with APOE4 homozygotes showing higher levels than heterozygotes.

**Figure 1:**
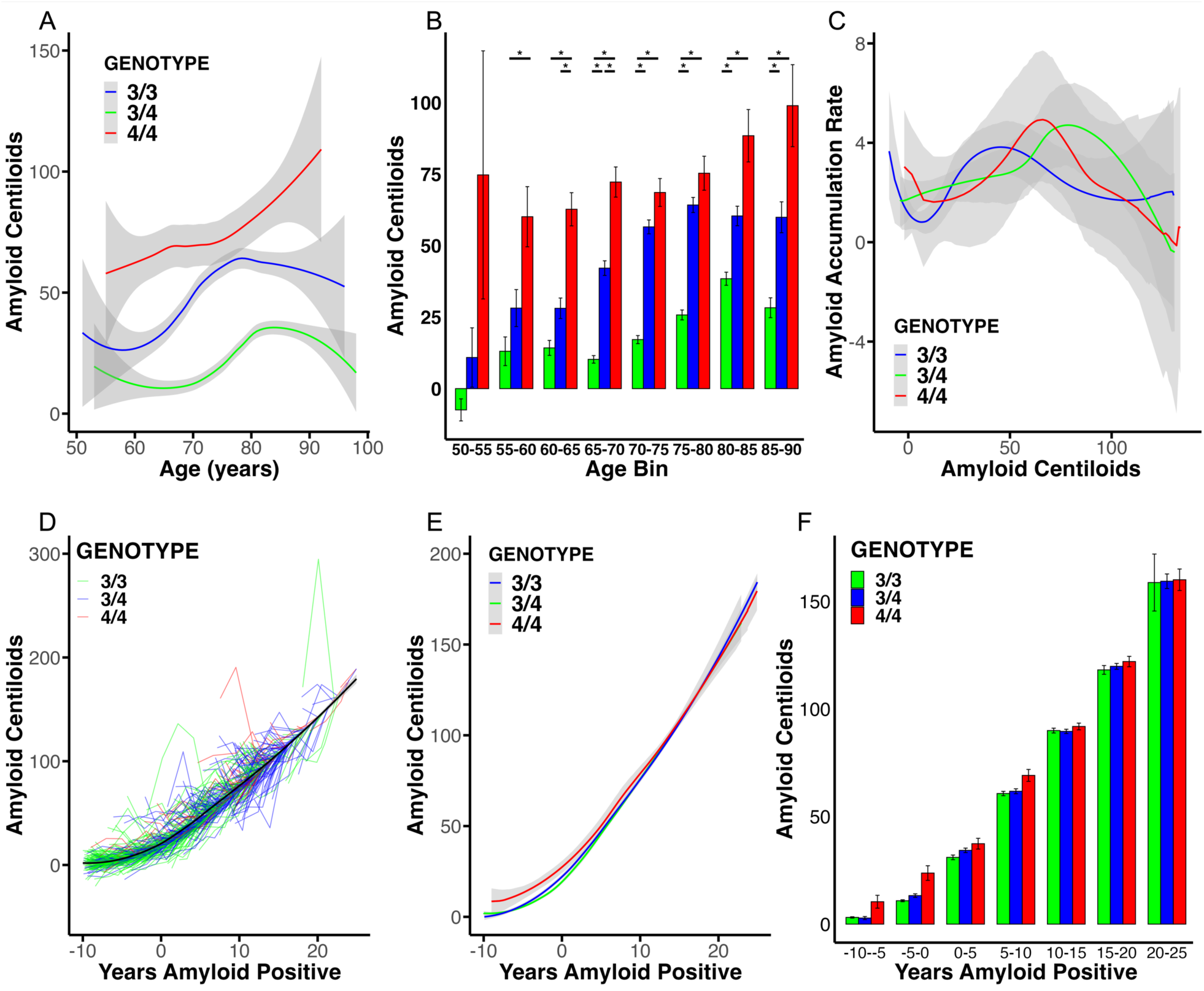
Aβ deposition as a function of APOE genotype. A) Aβ deposition increases with age, with APOE4/4 showing the highest levels, followed by APOE3/4 and APOE3/3. B) Binning by 5-year age blocks shows similar patterns, with differences emerging at age 55. C) Amyloid accumulation rate is similar across genotypes as a function of baseline amyloid. D) SILA model fit for estimated years of Aβ positivity across the dataset. E) SILA model fit split by genotype. F) No differences in Aβ deposition by estimated years of positivity after SILA modeling.

To further assess this, we binned individuals into five-year age groups (Figure 1B, and found significant effects of age group, APOE genotype, and an interaction (Two-way ANOVA; F_genotype_(2,3273)=355.26, p < 0.001; F_age bin_(7,3273)=30.83, p < 0.001; F_genotype×age bin_(14,3273)=3.29, p < 0.001). Post hoc tests showed no differences in Aβ levels between genotypes in individuals aged 50–55 (p > 0.87). However, from 55 to 60 years, significant differences emerged between APOE4/4 and APOE3/3 (p = 0.0064), with all genotypes diverging significantly in the 60–65 and 65–70-year age ranges (all p’s < 0.0001). Above age 70, APOE4 carriers (both homozygotes and heterozygotes) had significantly higher Aβ levels than APOE3 homozygotes (all p’s < 0.0001) but did not differ from each other (p’s > 0.12). These results suggest that APOE4 is associated with earlier Aβ deposition in a dose-dependent manner, with homozygotes accumulating Aβ earlier than heterozygotes, who in turn accumulate Aβ earlier than APOE3/3 carriers.

It is plausible that the dose-dependent increase in Aβ associated with the APOE4 genotype could be driven by a higher prevalence of Aβ positive individuals, therefore, we restricted analyses to Aβ positive participants. LOESS smoothing again showed increasing Aβ centiloid values across the age range. APOE3/4 and APOE4/4 genotypes were associated with significantly higher AUCs compared to APOE3/3, though the difference between APOE4/4 and APOE3/4 was not statistically significant (APOE3/3 [960, 1539], APOE3/4 [1742, 2310], APOE4/4 [2160, 2964]). Next, after splitting Aβ positive participants by age bin, we again found a significant effect of APOE genotype (F(2, 2432) = 158.45, p < 0.001), age group (F(7, 2432) = 21.91, p < 0.001), and a significant genotype-by-age interaction (F(13, 2432) = 3.54, p < 0.001). Post-hoc tests revealed no significant differences in Aβ deposition between genotypes before age 60. However, this result may reflect limited data, as there were fewer than 10 APOE4 homozygotes in this age range. From 60 to 65 years, APOE4 homozygotes had significantly higher Aβ levels compared to both APOE3/3 (p = 0.0007) and APOE3/4 (p = 0.009), with no difference between APOE3/3 and APOE3/4 (p = 0.999). In the 65 to 70 age range, APOE4 homozygotes exhibited higher Aβ levels compared to both APOE3/3 and APOE3/4 (p’s < 0.001) and APOE3/4 also showed elevated Aβ levels compared to APOE3/3 (p < 0.001). From 70 to 80 years, the same pattern was observed, although no difference was found between APOE4 homozygotes and heterozygotes (70-75: p = 0.99, 75-80: p = 0.99). In the 80 to 85 and 85 to 90 age groups, APOE4 homozygotes continued to have higher Aβ levels compared to APOE3/3 (p = 0.004, p = 0.01, respectively), though differences between APOE3/4 and either APOE3/3 or APOE4 homozygotes were not significant (p’s > 0.10). These findings further support the notion that APOE4 is associated with earlier Aβ deposition, even among Aβ individuals, with heterozygotes accumulating Aβ at younger ages than APOE3/3 individuals and homozygotes showing the earliest onset.

Given the association of APOE4 with increased Aβ burden across the age range, we hypothesized that APOE4 might also be linked to a faster rate of Aβ accumulation. To test this, we identified individuals with multiple Aβ PET scans and calculated their annual Aβ accumulation by taking the difference in cortical Aβ composite centiloids between their most recent and first scans, divided by the number of years between scans. We then plotted Aβ accumulation rates as a function of mean Aβ deposition (Figure 1C). Comparing the AUCs across genotypes, we found no significant difference in the rate of Aβ accumulation at different Aβ levels (APOE3/3 [218, 463], APOE3/4 [281, 474], APOE4/4 [70.7, 439]) These findings suggest that while APOE4 is associated with greater Aβ deposition, the rate of Aβ accumulation does not significantly differ between genotypes once an individual becomes Aβ positive.

Next, we graphed Aβ centiloid values as a function of age, with individual lines representing each participant (Figure 1D). Applying SILA to model the data, we found a strong relationship between Aβ deposition and estimated Aβ duration. When fitting separate LOESS curves, we did not observe substantial differences by genotype with the AUCs of the curves not differing significantly (APOE3/3 [1392, 1477], APOE3/4 [1429, 1498], APOE4/4 [1441, 1626]) (Figure 1E). To further investigate, we divided Aβ centiloid values into five-year blocks and examined the relationship between Aβ duration and Aβ levels by genotype (Figure 1F). While we found significant main effects of genotype and Aβ duration, we did not identify a significant interaction between genotype and Aβ duration (Two-way ANOVA; F_Aβ year bin_(5)= 3331.94, p < 0.00001, F_genotype_(2)=1907.48, p < 0.00001, F_genotype× Aβ year bin_ (10)=1.06, p = 0.39). This suggests that after normalizing the data to an Aβ threshold, there is no difference in Aβ deposition across genotypes.

### 3.2. Aβ duration is associated with selective loss of episodic memory

Given that APOE4 is linked to higher Aβ deposition but not a faster rate of accumulation, the reason APOE4 carriers experience greater memory loss remains unclear. We examined two potential explanations. First, APOE4 carriers may be further along the Aβ spectrum, with increased pathology driving memory decline. Alternatively, APOE4 may increase susceptibility to Aβ, such that less pathology would be needed to cause memory deficits in these individuals. To test these, we examined the relationship between estimated Aβ duration and several memory measures, including RAVLT delayed recall, RAVLT recognition, and delayed logical memory.

We began by generating LOESS curves for Aβ duration and RAVLT Delayed Recall (Figure 2A). Although the curves for each genotype began at similar points, APOE3/4 and APOE4/4 individuals experienced faster declines than APOE3/3. Comparing bootstrapped AUCs, we observed significantly lower AUCs for both APOE4 homozygotes and heterozygotes compared to APOE3 homozygotes, with a more pronounced decline in APOE4 homozygotes (APOE3/3 [– 3.31, 1.73], APOE3/4 [–11.0, –7.56], APOE4/4 [–26.6, –19.2]).

**Figure 2:**
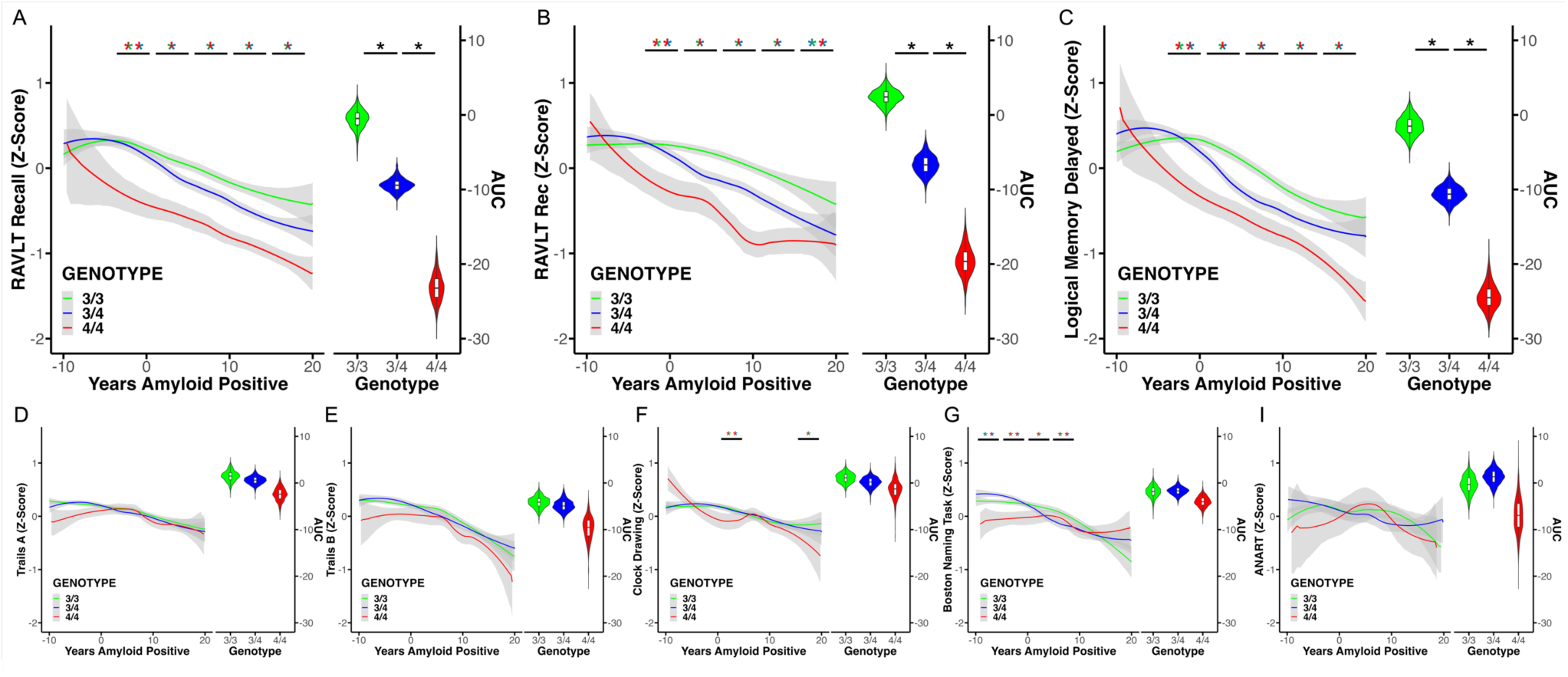
APOE4 accelerates episodic memory deficits as a function of amyloid duration. APOE4/4 is associated with faster decline in A) RAVLT delayed recall, B) RAVLT recognition, and C) Logical Memory delayed recall, compared to APOE3/4, which declines faster than APOE3/3. No significant differences in decline rates were observed between genotypes for D) TMT A, E) TMT B, F) Clock Drawing, G) BNT, and H) ANART. Confidence intervals for LOESS smooths were generated via bootstrapping (*n* = 1000), with violin plots displaying AUCs of bootstrapped lines. Significance derived from binned analyses (colors of * indicate comparisons): 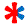 indicates a significant difference between APOE3/3 and APOE4/4; 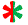 between APOE3/3 and APOE3/4; and 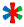 among all genotypes.

Next, we grouped participants into five-year Aβ duration blocks to determine when performance began to diverge across APOE genotypes. A significant main effect of Aβ duration (Supplemental Figure 1A, F(6,6285) = 86.84, p < 0.001), genotype (F(2,6285) = 240.37, p < 0.001), and a genotype-by-Aβ duration interaction (F(12,6285) = 2.49, p = 0.003) revealed that the relationship between Aβ positivity and cognitive decline varied by genotype. Post-hoc analyses indicated no differences in RAVLT delayed recall between genotypes in individuals five to 10 years before Aβ onset (p’s > 0.19). However, during the five years before Aβ onset, APOE4 homozygotes performed significantly worse than both APOE3 homozygotes (p < 0.001) and APOE4 heterozygotes (p < 0.001), while performance between APOE4 heterozygotes and APOE3/3 individuals did not differ (p = 0.99). Once individuals were Aβ positive, both APOE3/4 and APOE4/4 individuals exhibited worsening performance up to 20 years, with a more severe impairment in APOE4/4 compared to APOE3/4 (all p’s < 0.01). In the final 20–25– year block, APOE4 homozygotes performed worse than the other two groups (APOE3/3: p = 0.003; APOE3/4: p = 0.023), and no reliable difference was observed between APOE3/4 and APOE3/3 (p = 0.96). These results suggest that while memory decline accompanies increasing Aβ accumulation across all genotypes, APOE4 carriers exhibit a more rapid and severe decline, with APOE4 homozygotes experiencing the earliest and most significant decline.

We next examined whether recognition memory performance on the RAVLT followed a similar pattern to delayed recall (Figure 2B). Similar to delayed recall, recognition memory declined with longer Aβ duration, with APOE4 homozygotes and heterozygotes showing significantly lower AUCs for recognition memory compared to APOE3 homozygotes, and the decline was more pronounced in APOE4 homozygotes (APOE3/3 [0.34, 4.44], APOE3/4 [-9.05, –4.06], APOE4/4 [-23.3, –15.7]). We then grouped participants into five-year estimated Aβ duration blocks and observed significant main effects of Aβ duration (Supplemental Figure 1B, F(6,6163) = 64.70, p < 0.001), genotype (F(2,6163) = 243.89, p < 0.001), and a significant interaction between genotype and Aβ duration (F(12,6163) = 5.60, p < 0.001). Post-hoc analyses revealed no significant differences in recognition memory performance between genotypes five to ten years before the onset of Aβ positivity (all p’s > 0.10). Interestingly, however, during the five years before Aβ positivity, APOE4 homozygotes performed reliably worse than heterozygotes and APOE3/3 individuals, (p’s < 0.001), while heterozygotes did not reliably differ from APOE3/3 individuals (p = 0.25). Starting at the onset of Aβ positivity up to 15 years of Aβ positivity, APOE3/4 and APOE4/4 individuals demonstrated lower recognition memory compared to APOE3/3, with APOE4/4 individuals performing even worse than APOE3/4 (all p’s < 0.01). During the 15-to-20-year positivity bin, we found that APOE4 homozygotes and heterozygotes performed worse than APOE3/3 individuals (p’s < 0.02) but did not differ from each other (p = 0.17). Further, during the 20-to-25-year block, APOE4/4 individuals showed worse performance than APOE3/3 (p = 0.01) but not APOE3/4 individuals (p = 0.19), and there was no difference between APOE3/4 and APOE3/3 (p = 0.11). These results again suggest that recognition memory declines earlier in APOE4 carriers and is more severe in APOE4 homozygotes.

Lastly, we examined memory ability on a separate task, the Logical Memory test, finding that performance on this test decreased with increased Aβ duration and that this decline was exacerbated in APOE4 carriers (Figure 2C). Comparing the bootstrapped AUCs, we again found deceased AUCs for both APOE3/4 and APOE4/4 individuals compared to APOE3/3, and APOE4/4 was associated with lower memory scores compared to APOE3/4 (APOE3/3 [-3.94, 1.06], APOE3/4 [-12.7, –8.55], APOE4/4 [-27.5, –20.6]). When we categorized individuals based on how long they were estimated to have been Aβ positive, we found significant effects for Aβ duration (Supplemental Figure 1C, F(6,5489) = 118.05, p < 0.001), genotype (F(2,5489) = 239.99, p < 0.001), and their interaction (F(12,5489) = 5.13, p < 0.001). The post-hoc results mirrored what we observed with the RAVLT. We found that APOE4 heterozygotes performed better than APOE3/3 individuals (p = 0.01) while APOE4 homozygotes did not differ from other genotypes (p’s > 0.65). Conversely, APOE4 homozygotes who were within five years of Aβ positivity performed reliably worse than APOE4 heterozygotes and APOE3 homozygotes with no differences between the latter two genotypes (3/3 vs 3/4: p = 0.63; 3/3 vs 4/4: p < 0.001; 3/4 vs 4/4: p < 0.001). Once individuals were Aβ positive, APOE4 homozygotes performed reliably worse up to twenty years post positivity compared to both APOE3/3 and APOE3/4 individuals (all p’s < 0.01) and APOE3/4 individuals performed reliably worse than APOE3/3 individuals (p’s < 0.05). During the 20-to-25-year block, APOE4 homozygotes performed reliably worse that APOE3/3 individuals (p = 0.003), marginally worse than APOE3/4 individuals (p = 0.05) with APOE3/3 and APOE3/4 individuals not reliably differing in performance (p = 0.19). These results suggest that APOE4 may increase susceptibility to Aβ, such that less Aβ is required to trigger memory decline in carriers compared to non-carriers.

### 3.3. APOE4 carriers do not exhibit increased decline on non-memory tasks

Given that APOE4 carriers exhibited accelerated memory loss in the presence of Aβ, we next explored whether this effect is specific to the memory domain or occurs across other cognitive domains as well. To address this, we examined performance on the TMT A and B, the Clock Drawing Task, the BNT, and the ANART, analyzing how these abilities changed as a function of estimated years of Aβ positivity (Figure 2D-I).

Across non-memory domains, we did not observe clear differences between APOE genotypes. While all domains declined as a function of amyloid duration, the rate and extent of decline did not systematically vary by genotype. Specifically, when comparing bootstrapped AUCs, no reliable differences emerged between genotypes on any task (Table 2). To further investigate, we binned performance into five-year intervals and found no significant interaction between amyloid duration and APOE genotype for the TMT A, TMT B, or the ANART (Supplemental Figure 1D-I, Two-way ANOVA interaction terms, p’s > 0.12). However, we did observe significant interactions between amyloid duration and APOE genotype for the Clock Drawing Task (Figure 3F, Supplemental Figure 1F) and the BNT (Figure 3G, Supplemental Figure 1G).

**Figure 3:**
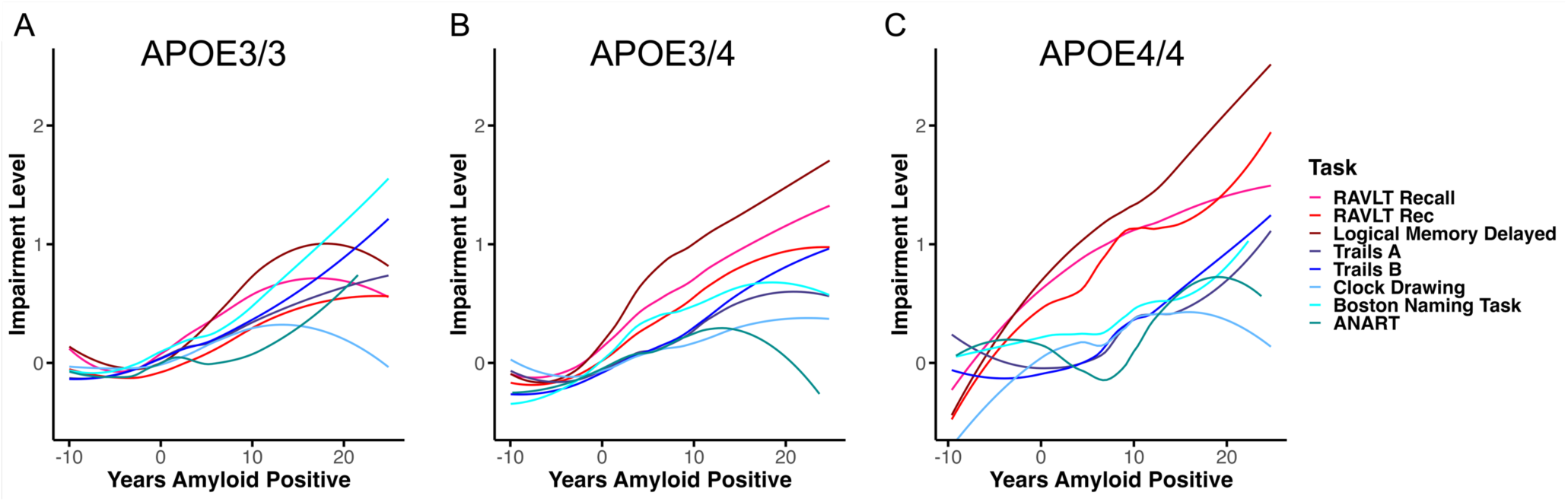
Patterns of cognitive impairment by estimated years of Aβ positivity. A) In APOE3/3, episodic memory impairments emerge approximately seven years after Aβ positivity, followed shortly by deficits in other domains. B) In APOE3/4, episodic memory impairment begins five years after Aβ positivity onset, with other domains affected thereafter. C) In APOE4/4, episodic memory impairment appears at or before Aβ positivity onset, while impairments in other domains emerge nearly a decade later. Memory tasks in hot colors and non-memory tasks in cool colors.

**Table 2:** Performance on non-memory tasks as a function of APOE genotype.

| Task | APOE3/3 | APOE3/4 | APOE4/4 |
| --- | --- | --- | --- |
| **TMT A** | [-0.870 3.71] | [-1.05 2.32] | [-5.38 0.288] |
| **TMT B** | [-6.61 -1.96] | [-7.16 -2.26] | [-14.9 -5.21] |
| **Clock Drawing Task** | [-0.992 3.08] | [-2.08 2.06] | [-5.77 2.37] |
| **BNT** | [-4.62 0.381] | [-3.47 0.0519] | [-6.24 -1.98] |
| **ANART** | [-4.42 2.98] | [-1.86 4.40] | [-13.9 0.659] |

Table 2: Performance on non-memory tasks as a function of APOE genotype
| **Task** | **APOE3/3** | **APOE3/4** | **APOE4/4** |
| --- | --- | --- | --- |
| **TMT A** | [-0.870 3.71] | [-1.05 2.32] | [-5.38 0.288] |
| **TMT B** | [-6.61 -1.96] | [-7.16 -2.26] | [-14.9 -5.21] |
| **CDT** | [-0.992 3.08] | [-2.08 2.06] | [-5.77 2.37] |
| **BNT** | [-4.62 0.381] | [-3.47<br>0.0519] | [-6.24 -1.98] |
| **ANART** | [-4.42 2.98] | [-1.86 4.40] | [-13.9 0.659] |

For the Clock Drawing Task, we found significant main effects of Aβ duration (F(6,6121) = 28.71, p < 0.001), genotype (F(2,6121) = 18.73, p < 0.001), and a genotype-by-Aβ duration interaction (F(12,6121) = 1.89, p = 0.03). We found that APO4 homozygotes performed worse than both heterozygotes and APOE3 homozygotes during the first five years of amyloid positivity (p’s < 0.05), while the latter two genotypes did not differ (p = 0.99). The only other reliable difference was observed at the 15-to-20-year block with APOE4 homozygotes performing worse than APOE3 homozygotes (p = 0.04). All other comparisons between groups and across year bins were not reliably different (all p’s > 0.13). For the BNT, we found significant main effects of Aβ duration (F(6,4180) = 57.56, p < 0.001) and genotype (F(2,4180) = 45.04, p < 0.001), as well as a significant genotype-by-Aβ duration interaction (F(12,4180) = 3.85, p < 0.001). However, post-hoc analyses indicated no significant differences between genotypes at most time points, suggesting that despite the interaction, the observed effects were not strong enough to differentiate between groups. We found that APOE4 heterozygotes performed better than both APOE3 homozygotes (p = 0.04) and APOE4 homozygotes (p = 0.01) while APOE3/3 and APO4/4 did not significantly differ (p = 0.08). We also found that APOE4 homozygotes performed significantly worse than both APOE3/3 and APOE3/4 individuals at the 0-to-5-year block (APOE3/3 vs APO3/4: p = 0.99, APOE3/3 vs APO4/4: p = 0.006, APOE3/4 vs APO4/4: p = 0.008). However, APOE4 homozygotes did not exhibit impairment compared to the other APOE genotypes on any of the other blocks. We also found that APOE3/4 individuals performed marginally better compared to APOE3/3 individuals five years before positivity to the threshold (p = 0.050). while they performed worse at the 5-to-10-year bin (p = 0.001). We found no other significant differences between APOE genotypes (all p’s > 0.05). Overall, the results show that the decline of the Clock Drawing Task and BNT does not systematically vary between APOE3/3, APOE3/4, and APOE4/4 carriers.

### 3.4. No Reliable differences by Sex

Research has highlighted the critical role of sex in modulating APOE4-related AD pathology. In our primary analysis, we controlled for sex effects; however, in a post-hoc analysis, we explored whether the accelerated episodic memory decline observed in APOE carriers was influenced by sex. Using LOESS smoothing with bootstrapping, we stratified the data by genotype and sex and calculated AUCs from the bootstrapped results. Our findings revealed that males consistently performed worse than females across all three episodic memory measures: RAVLT Delayed Recall, RAVLT Recognition, and Logical Memory Delayed Recall. However, these performance differences did not vary reliably by APOE genotype (Supplemental Table 1). These results align with prior research showing that males generally exhibit poorer episodic memory performance compared to females These results largely recapitulate prior work demonstrating that males perform worse on episodic memory compared to females (Bleecker et al., 1988; Gale et al., 2007). However, we found that sex did not significantly moderate episodic memory decline within APOE genotypes.

### 3.5. Cognitive trajectories in AD differ by APOE genotypes

Given the substantial differences in how APOE genotypes affect various cognitive domains, we aimed to understand the time course of cognitive changes as a function of APOE genotype. To do this, we derived impairment values for all Aβ positive individuals (Figure 4). Consistent with prior work, we z-scored all individuals relative to Aβ negative individuals and fit LOESS curves for each cognitive measure as a function of Aβ duration (Li et al., 2024). We first examined the cognitive trajectory for APOE3/3 individuals. Initially, the memory tasks, RAVLT delayed recall and logical memory, were the first to deviate from Aβ negative performance after around 7 years of Aβ positivity. However, this was soon followed by impairments across other cognitive domains. For APOE3/4 individuals, all three memory measures began to show impairment after around 5 years of Aβ positivity, with impairments in the non-memory domains emerging a few years later.

In contrast, for APOE4/4 individuals, memory impairments were severe, potentially starting at or even before Aβ positivity and progressing linearly, reaching a much higher level of impairment compared to both APOE3/3 and APOE3/4 individuals. Notably, non-memory impairments only began after around 12 years of Aβ positivity and memory impairment had begun. Despite differences in timing and severity, the pattern of cognitive decline was consistent across genotypes: memory declines emerged first, with APOE4 being associated with earlier and more severe memory impairments but not greater impairments in other cognitive domains.

## 4. Discussion

In this study, we investigated whether APOE4 carriers were more susceptible to Aβ accumulation than non-carriers. Specifically, we examined whether cognitive trajectories diverged among APOE3/3, APOE3/4, and APOE4/4 carriers across multiple cognitive domains, including episodic memory, executive function, processing speed, and language ability. We found that APOE4 carrier status was selectively associated with an accelerated cognitive decline in the presence of Aβ. APOE4 carriers began to show episodic memory decline after fewer years of Aβ positivity and declined more severely compared to APOE3/3 individuals. Additionally, there was a dose-dependent effect, with individuals carrying two copies of APOE4 exhibiting even greater impairment than those with only one copy. Importantly, this decline was selective to episodic memory, and we did not observe a similar pattern in the other cognitive domains. The earlier and more severe memory decline in these individuals suggests that interventions targeting Aβ deposition may need to be initiated earlier in APOE4 carriers to slow or prevent the onset of cognitive impairment.

### 4.1. Selective decline in episodic memory in APOE4 carriers

APOE4 carriers exhibit larger deficits in episodic memory with increasing age and decline faster in this domain over time (Bondi et al., 1995; Eich et al., 2019; Sinha et al., 2018). However, research suggests that this decline is driven by APOE4 carriers with elevated Aβ, as older adults with one or two copies of APOE4 but without elevated Aβ did not exhibit longitudinal episodic memory decline. While it is well known that Aβ deposition is linked to episodic memory decline, these findings suggest that APOE4 may exacerbate this effect. However, it remains unclear whether APOE4 carriers experience faster memory decline because they accumulate more Aβ (less resistance) or because they are more vulnerable to the effects of Aβ, requiring less Aβ to trigger cognitive decline (less resilience).

Given that APOE4 carriers experience an earlier onset of Aβ deposition but do not accumulate Aβ at a faster rate once positive, we used SILA to estimate the year a person becomes Aβ positive (Betthauser et al., 2022; Fortea et al., 2024; Lim et al., 2017). Using these estimates, we quantified cognitive performance across a comprehensive battery of neuropsychological assessments as a function of Aβ positivity duration. We demonstrated that APOE3/4 and APOE4/4 individuals exhibit episodic memory deficits after being Aβ positive for a shorter period compared to APOE3/3 individuals. Moreover, deficits remained more pronounced in APOE carriers even after being Aβ positive for 25 years. Additionally, this decline was dose-dependent, with individuals carrying two copies of APOE4 exhibiting even more severe memory impairment compared to those with only one copy. Importantly, this decline was specific to episodic memory and was not observed in other cognitive domains, indicating that the effect is not simply a result of overall faster cognitive decline but rather a memory-specific vulnerability.

Interestingly, we did not observe significant sex effects in this study. Specifically, the accelerated episodic memory decline in APOE4 carriers was not selective to males or females. Prior work has suggested that sex may mediate the effects of APOE4 on cognitive function (Williams et al., 2019). However, these results have been contradictory with studies finding more rapid decline in males, while others find faster decline in females (Buckley et al., 2018; Wang et al., 2019). Importantly, these studies either did not focus on Aβ deposition or did not split performance across different cognitive domains.

### 4.2. Potential mechanisms of episodic memory decline

Episodic memory critically relies on the hippocampus, which is one of the earliest regions affected, both directly and indirectly (e.g., degrading its input via damage to the entorhinal cortex), in the pathogenesis of AD. In fact, age-related and AD-related changes within the hippocampus are strong predictors of episodic memory deficits (Adams et al., 2023; de Leon et al., 1996; Radhakrishnan et al., 2022). Moreover, APOE4 has been associated with numerous pathological changes in the hippocampus, including synapse loss, atrophy, blood-brain barrier degradation, and alterations in activity patterns (Fernández-Calle et al., 2022; Montagne et al., 2020; Najm et al., 2019; Snellman et al., 2023). Specifically, APOE4 carriers exhibit more rapid hippocampal atrophy over time, and the link between hippocampal dysfunction and episodic memory decline is stronger in carriers compared to non-carriers (Håglin et al., 2023).

One potential mechanism underlying this vulnerability arises from altered hippocampal activity in APOE4 carriers. Research has shown that APOE4 carriers exhibit increased hippocampal activity, with this hyperactivity detectable as early as young adulthood (Filippini et al., 2009). Notably, hippocampal hyperactivity has been linked to both increased Aβ deposition and tau accumulation (Adams et al., 2022; Giorgio et al., 2024; Leal et al., 2017). Studies suggest that Aβ deposition may drive this hyperactivity, which in turn may promote tau accumulation and its spread beyond the medial temporal lobe which is a significant risk factor for cognitive impairment and dementia (Adams et al., 2022; Corriveau-Lecavalier et al., 2024; Giorgio et al., 2024).

Therefore, APOE4 may preferentially alter hippocampal function, resulting in more rapid hippocampal dysfunction. This could explain why APOE4 carriers exhibit more rapid and severe memory impairment, and why this decline is selective to memory rather than other cognitive domains that do not highly dependent on hippocampal function. Additionally, hippocampal hyperactivity may also contribute to Aβ deposition. Research has shown that while Aβ deposition is linked to increased neural activity, there is also evidence suggesting a reverse relationship, where increased activation promotes Aβ deposition (Giorgio et al., 2024; Leal et al., 2017; Oh et al., 2015). If this pattern is cyclical, APOE4 carriers may be at higher risk for Aβ accumulation, potentially explaining why these individuals develop Aβ at younger ages. This increased activation may also contribute to the more rapid episodic memory decline observed in APOE4 carriers.

It is important to note that there are many other potential mechanisms that may contribute to the cognitive changes observed in APOE4 individuals. For instance, hippocampal neuroinflammation is an early pathology in AD, and it is plausible that APOE4 may exacerbate this process (Calsolaro & Edison, 2016; Janelidze et al., 2018; Noche et al., 2024). Additionally, there may be other pathologies initiated by APOE4 that have yet to be identified, which could contribute to hippocampal dysfunction. Understanding the mechanisms that drive the accelerated memory decline in APOE4 carriers is crucial for identifying biomarkers and developing novel therapeutic targets to treat this vulnerable population.

### 4.3. Implications

It is widely believed that cognitive decline lags behind Aβ deposition by up to three decades (Jack et al., 2018; Sperling et al., 2013). Our work demonstrates that, while cognitive decline does indeed follow Aβ deposition, the timeline varies by APOE genotype and cognitive domain. Therefore, when diagnosing and monitoring AD, it is crucial to recognize that cognitive changes are not uniform across the population and may be influenced by APOE status. Significant research has focused on using subtle memory changes to identify individuals with elevated AD biomarkers and a high risk of future decline (Holmqvist et al., 2023; Thomas et al., 2018, 2020; Vanderlip, Stark, et al., 2024). In this context, APOE4 carriers may be an especially suitable population for these tools, as they tend to exhibit memory deficits earlier in disease progression. However, this also suggests that, when possible, cognitive assessments should be interpreted in conjunction with APOE4 genotype, and the criteria for identifying impairment on these tasks may need to differ based on APOE status.

Another important question is whether Aβ positivity thresholds should vary across genetic groups, such as APOE4 homozygotes. Classifying individuals as Aβ-positive or Aβ-negative helps identify those experiencing pathological changes consistent with Alzheimer’s disease (Jack et al., 2024; Jagust et al., 2021; Jansen et al., 2022). However, it is possible that APOE4 carriers exhibit lower Aβ levels, meaning the current threshold may not capture individuals at comparable stages of disease progression. In this study, we used a lower threshold of 16.7 centiloids compared to other research, but even this may be too high for APOE4 carriers. An even lower threshold could allow for earlier diagnoses, interventions, and treatment strategies in this group. However, this may not be currently possible with current PET tracers and quantification techniques.

An interesting finding that has emerged is that APOE4 carriers tend to derive reduced benefits from therapies targeting AD pathology. For instance, anti-Aβ therapies, such as lecanemab, do not appear to significantly slow cognitive decline in APOE4 carriers, and APOE4 homozygotes on these therapies have even shown a qualitative exacerbation of cognitive decline compared to those on placebo (Van Dyck et al., 2023). Additionally, a recent study using low-dose levetiracetam to reduce hippocampal hyperactivity in AD found benefits only in APOE4 non-carriers, with no improvements observed in carriers (Mohs et al., 2024). The lack of efficacy of these therapies in APOE4 carriers may be because these individuals have already experienced substantial pathological changes, such as hippocampal atrophy or cortical tau accumulation. Moreover, APOE4 carriers may harbor additional, yet unidentified, pathologies that need to be addressed to effectively slow cognitive decline in this population.

## 5. Conclusion

In this study, we demonstrated that APOE4 carriers are particularly vulnerable to Aβ pathology, exhibiting accelerated episodic memory decline along the AD spectrum. Furthermore, this decline followed a dose-dependent pattern, with APOE4 homozygotes showing the fastest and most severe impairment. These findings suggest that the time course and magnitude of cognitive decline in AD may vary depending on APOE genotype. It is essential to determine whether APOE4 carriers experience fundamentally different disease progression and whether distinct biomarkers and treatment approaches are required to prevent and treat cognitive decline in this population.

## 6. Acknowledgements

Data collection and sharing for this project was funded by the Alzheimer’s Disease Neuroimaging Initiative (ADNI) (National Institutes of Health Grant U01 AG024904) and DOD ADNI (Department of Defense award number W81XWH-12-2-0012). ADNI is funded by the National Institute on Aging, the National Institute of Biomedical Imaging and Bioengineering, and through generous contributions from the following: AbbVie, Alzheimer’s Association; Alzheimer’s Drug Discovery Foundation; Araclon Biotech; BioClinica, Inc.; Biogen; Bristol-Myers Squibb Company; CereSpir, Inc.; Cogstate; Eisai Inc.; Elan Pharmaceuticals, Inc.; Eli Lilly and Company; EuroImmun; F. Hoffmann-La Roche Ltd and its affiliated company Genentech, Inc.; Fujirebio; GE Healthcare; IXICO Ltd.; Janssen Alzheimer Immunotherapy Research & Development, LLC.; Johnson & Johnson Pharmaceutical Research & Development LLC.; Lumosity; Lundbeck; Merck & Co., Inc.; Meso Scale Diagnostics, LLC.; NeuroRx Research; Neurotrack Technologies; Novartis Pharmaceuticals Corporation; Pfizer Inc.; Piramal Imaging; Servier; Takeda Pharmaceutical Company; and Transition Therapeutics. The Canadian Institutes of Health Research is providing funds to support ADNI clinical sites in Canada. Private sector contributions are facilitated by the Foundation for the National Institutes of Health (www.fnih.org). The grantee organization is the Northern California Institute for Research and Education, and the study is coordinated by the Alzheimer’s Therapeutic Research Institute at the University of Southern California. ADNI data are disseminated by the Laboratory for Neuro Imaging at the University of Southern California.

## 7. Funding

This research was funded in part by R01 AG066683 (CS) and P30 AG066519 (CS).

## 8. Competing interests

The authors report no competing interests.

## Notes

### Competing Interest Statement

The authors have declared no competing interest.

