## Supplemental Information for "APOE4 Increases Susceptibility to Amyloid, Accelerating Episodic Memory Decline"

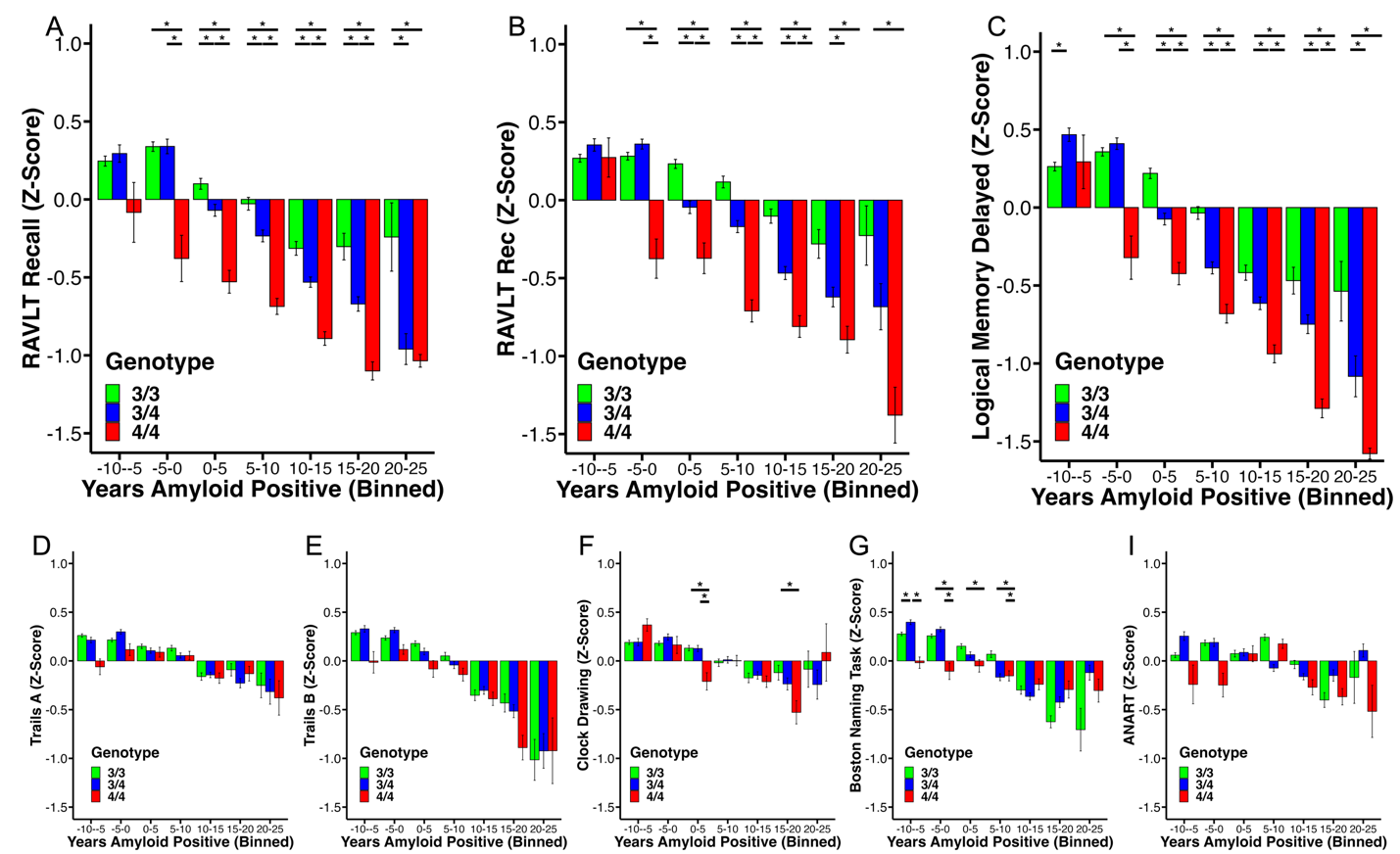

**Supplemental Figure 1: Neuropsychological performance split by five-year bins of estimated years of Aβ positivity.** APOE4/4 shows significant impairment on RAVLT delayed recall, RAVLT recognition, and Logical Memory delayed recall relative to APOE3/4 and APOE3/3, beginning five years before amyloid positivity. APOE3/4 also shows impairment relative to APOE3/3 beginning at onset of Aβ positivity. Increased years of Aβ positivity is associated with decline on D) Trails A and E) Trails B, with no genotype differences. F) APOE4/4 shows worse performance on the Clock Drawing Task during the first 5 years and between 15–20 years of Aβ positivity. G) APOE4/4 has reduced performance on the Boston Naming Task before amyloid positivity and up to 10 years post-onset, with no differences in later periods. I) ANART performance shows no differences by APOE genotype. Bar graphs represent mean ± SEM.

**Supplemental Table 1:**

| **Task** | **All** | | **APOE3/3** | | **APOE3/4** | | **APOE4/4** | |
| --- | --- | --- | --- | --- | --- | --- | --- | --- |
|  | Male | Female | Male | Female | Male | Female | Male | Female |
| **RAVLT Delayed Recall** | [-14.2. -10.9] | [-4.67 -1.07] | [-8.83 -1.96] | [2.93 8.06] | [-15.4 -10.5] | [-5.78 -1.11] | [-30.8 -22.8] | [-24.9 -15.6] |
| **RAVLT Recognition** | [-8.82 -4.40] | [-7.04 -2.71] | [-3.56 1.82] | [2.97 8.72] | [-8.10 -2.74] | [-9.04 -3.55] | [-27.6 -14.5] | [-25.1 -16.2] |
| **Logical Memory Delayed Recall** | [-14.5 -10.2] | [-8.03. -3.76] | [-7.81 -2.21] | [0.190 5.37] | [-15.1 -9.52] | [-9.48 -3.71] | [-30.9 -19.1] | [-28.2 -19.2] |
